# *C. elegans* CED-3 caspase promotes non-apoptotic linker cell dismantling but not death onset

**DOI:** 10.1101/2025.10.15.682583

**Authors:** Olya Yarychkivska, Lena M. Kutscher, Betty Ortiz Bido, Juliana Kagan, Yoon A. Kim, Dana Mamriev, Yun Lu, Elli Puscher, Wolfgang Keil, Sarit Larisch, Shai Shaham

## Abstract

Nuclear degradation accompanies cell death. To study this process, we followed nuclear dismantling of the *C. elegans* linker cell, which undergoes non-apoptotic morphologically-conserved cell death characterized by nuclear envelope crenellations and cell splitting. We show that although linker cell death is cell autonomous, nucleus elimination follows engulfment and is blocked in *rab-35* and *arf-6* phagosome maturation mutants. Surprisingly, although linker cell death is independent of the apoptotic caspase CED-3, CED-3 is partially required within the linker cell, and upstream of RAB-35 and ARF-6, for cell splitting, engulfment, and nucleus elimination. In parallel studies, we found that the kinase inhibitor staurosporine causes mouse embryonic fibroblasts to undergo caspase-independent non-apoptotic death accompanied by nuclear crenellations and, paradoxically, by *Caspase-3/7* activation. Our findings suggest mechanistic similarities between staurosporine-induced and linker cell death, revealing that, in some contexts, caspases do not initiate cell death but instead promote subcellular tasks required for cell clearance.

## Introduction

Programmed cell death is a complex process that effects the destruction of cellular compartments and their building blocks [1-3]. How specific subcellular structures and organelles are destroyed during cell death is not well understood. Some aspects of cellular destruction appear to be cell autonomous. For example, during apoptosis, caspases acting within dying cells cleave regulatory proteins, leading to DNAse activation and subsequent chromatin dismantling and DNA degradation [4]. By contrast, caspase-dependent cleavage of Xk-family proteins results in presentation of cell surface lipids, such as phosphatidyl serine, attracting phagocytes that engulf the dying cell and promote its non-autonomous degradation [5, 6]. Once engulfed, the dying cell resides in a phagosome, which then fuses with lysosomes [2, 7]. Phagolysosomes in some contexts can tubulate into smaller vesicles whose subsequent degradation is more facile. In the degradation of *C. elegans* polar bodies, for example, tubulation requires the small GTPase ARL-8 acting in the engulfing cell, and depends on amino acid export, activation of the lysosomal mTOR complex, mTORC, and subsequent recruitment of a BLOC-1-related complex [8]. Although it has been suggested that dying cell membrane breakdown is independent of autophagy [9], how dying cell organelles are dismantled within phagolysosomes has not been extensively investigated.

Cell death is a common cell fate during the development of the nematode *C. elegans*, with 131 of the 1090 cells born in the hermaphrodite undergoing this process [10, 11]. While nearly all of these cell death events are blocked by inhibiting the apoptotic caspase CED-3 [12, 13], the death of the male-specific linker cell is an exception [14]. The linker cell guides migration of the developing male gonad, and its death, at the larva-to-adult transition, facilitates fusion between the *vas deferens* and the cloaca, generating an open reproductive system [15, 16] (Figure 1A). Linker cell death is non-apoptotic by morphological and genetic criteria [14, 17]. Unlike apoptotic cells, the dying linker cell fails to exhibit chromatin condensation. Indeed, even nuclear periphery-associated heterochromatin disappears during early stages of linker cell death [14]. While the nuclear envelope of apoptotic cells remains smooth as cell demise proceeds, the dying linker cell nuclear envelope is characterized by nuclear crenellations, deep invaginations that can be detected by both light and electron microscopy (Figure 1A). Dying linker cells also exhibit swollen organelles, a feature reserved for only very late stages of apoptosis [3, 14, 18, 19].

**Figure 1.**
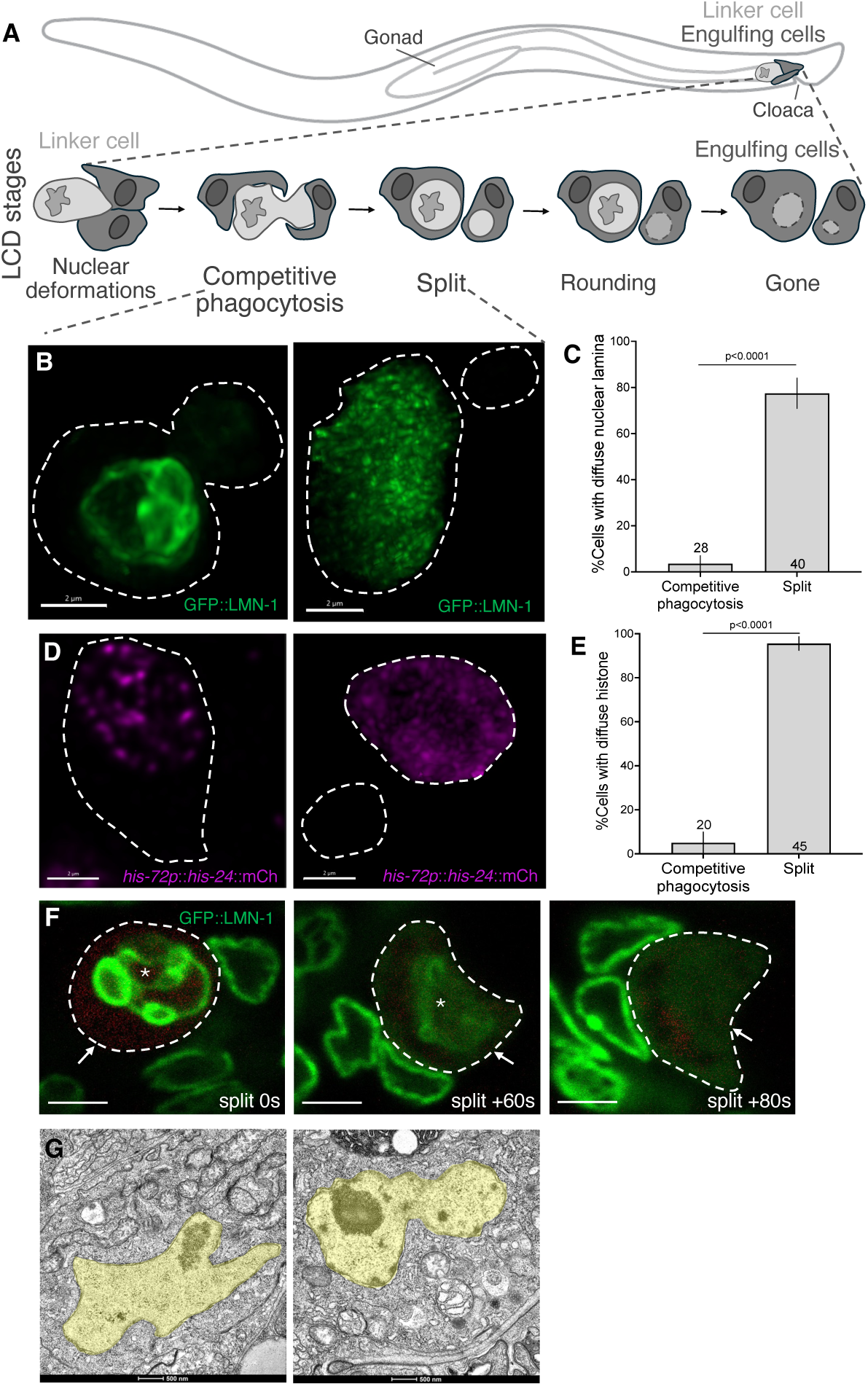
Nuclear dismantling initiates after linker cell splitting. (A) Linker cell death stages after the cell arrives at the tail. (B) Representative confocal images of endogenous reporter for lamin during of competitive phagocytosis (left) and splitting (right). Scale bar, 2μm. (C) Quantification of cells with diffuse lamin signal at indicated stages. Number of animals scored inside bars. Average of 3 independent experiments. Error bars, standard error of sample proportion, Fisher’s exact test. (D) Representative confocal images of HIS-24 (histone H1) during the stages of competitive phagocytosis (left) and splitting (right). Scale bar, 2μm. (E) Quantification of cells with diffuse histone signal at two stages. The number of animals scored inside bars. Average of 3 independent experiments. Error bars, standard error of sample proportion, Fisher’s exact test. (F) Long-term confocal imaging of GFP::LMN-1 in a single animal showing rapid lamina redistribution. Scale bar, 2μm. (G) Electron micrographs of linker cells at a competitive phagocytosis (left) and split (right) stages. Nucleus pseudocolored yellow. Scale bar, 500nm.

Genetically, linker cell death is independent of the main apoptotic caspase, CED-3, and of the three other caspase-related proteins encoded in the *C. elegans* genome [14, 17]. Linker cell death is also independent of other apoptotic effectors, including CED-4/Apaf1, CED-9/Bcl2, and EGL-1/BH3-only [14]. Instead, studies from our lab demonstrated that multiple upstream regulatory signals [14, 18-20] converge on the HSF-1 heat-shock factor, a transcriptional regulator that acts non-canonically to promote cell death instead of protecting against it [18]. During linker cell death, HSF-1 activates expression of target genes unrelated to the heat-shock response, including genes encoding the LET-70/UBE2D2 E2 ubiquitin ligase and other ubiquitin proteasome system components [18].

We previously showed that the linker cell undergoes stereotypical morphological changes as it dies [16]. Nuclear crenellation is followed by asymmetric cell splitting and subsequent engulfment of the resulting two fragments by different descendants of the U progenitor cell, U.I/rp (Figure 1A). Degradation of the linker cell nucleus-containing fragment is mediated by the RAB-35 and ARF-6 small GTPases, which prepare the phagosome in which the linker cell fragment resides for lysosomal fusion and dismantling [16].

To understand how cellular organelles are degraded during cell death, we studied the process of nuclear dismantling in linker cell death. We show, using a combination of live confocal imaging and electron microscopy, that disassembly of the nuclear lamina and displacement of histones from the nucleus takes place downstream of linker cell splitting and phagosome maturation, and before nuclear envelope degradation. Although linker cell death does not require engulfment, we find that in *rab-35(lf)* or *arf-6(gf)* mutants, nucleus disassembly stalls. Surprisingly, a similar blockade is evident in *ced-3(lf)* caspase and *ced-4(lf)* Apaf1 mutants. While cell death is initiated in these mutants, and the linker cell is eventually degraded in many of them, about half of adult males exhibit a persisting unengulfed cell corpse even 24 hours after cell death begins. We demonstrate that in *ced-3* mutants, cell splitting is defective, likely accounting for the engulfment defect. We also show that while the non-muscle myosin NMY-2 normally accumulates specifically in the smaller splitting fragment, it is more equally distributed between the two fragments in *ced-3* mutants.

Cell death induced by the non-specific kinase inhibitor staurosporine has been suggested to cause apoptosis; however, cell death ensues in cell lines established from the fetal thymus of mutants lacking key apoptotic effectors [21], suggesting a non-apoptotic mechanism. We demonstrate that staurosporine-induced death of mouse embryonic fibroblasts (MEFs) is independent of *Casp3* and *Casp7*. Importantly, fibroblast nuclei become crenellated, and expression of ubiquitin proteasome system components is induced, reminiscent of linker cell death.

Our studies suggest that back-and-forth signaling between the engulfing and dying cell may be necessary to trigger nuclear destruction during linker cell death; demonstrate a novel role for caspases in this cell death form; and suggest that linker cell-type death (LCD), in which caspases perform non-death auxiliary functions, may be conserved from *C. elegans* to mammals.

## Results

### Disruption of lamin and histone nuclear localization occurs after linker cell splitting and before nuclear membrane disassembly

To characterize nuclear dynamics during linker cell death, we imaged the localization of LMN-1 (the sole *C. elegans* nuclear lamin) endogenously tagged with GFP, as well as reporter transgenes for the nuclear proteins HIS-24 (Histone H1), EMR-1 (Emerin), and NPP-1 (NUP54), in animals at different stages of linker cell death. We found that while all these proteins are confined to the nucleus in animals in which the linker cell has not yet split, LMN-1, HIS-24, and NPP-1 take on a diffuse nucleocytoplasmic distribution in animals in which the linker cell has fragmented (Figure 1B-E, Supplemental Figure 1B). Localization of EMR-1 is less obviously disrupted, with at least some of the protein retaining what appears to be nuclear membrane association after linker cell splitting (Supplemental Figure 1A). None of these proteins are present in the small linker cell fragment generated after cell splitting.

To confirm these dynamics, we followed GFP::LMN-1 using a confocal long-term live-imaging microfluidic setup [22] in which linker cell death can be longitudinally followed in single animals. Corroborating our bulk time-course studies, we found that cell splitting precedes LMN-1 redistribution. The interval between these processes is variable (1 min to 3 h, n=7), but once initiated, LMN-1 redistribution occurs rapidly, within 60-80 seconds (Figure 1F). GFP::LMN-1 dissociation coincides with nuclear envelope blebbing visible by brightfield imaging (Supplemental Figure 1C), which can serve as a proxy for lamina dismantling. Electron microscopy of a split cell undergoing nuclear blebbing reveals that nuclear membranes are intact despite lamina redistribution (Figure 1G), consistent with partial retention of EMR-1 nuclear envelope localization (Supplemental Figure 1A).

Taken together, these studies demonstrate that nuclear dismantling during linker cell death is a multi-step process in which partial nuclear lamina disassembly and chromatin disruption follow cell splitting, while nuclear membranes are degraded only later.

### Lamin disassembly during linker cell death requires phagosome sealing

Previous work showed that linker cell splitting coincides with engulfment of the two linker cell fragments by the two U.l/rp cells, a process we have termed competitive phagocytosis [16]. To assess the temporal relationship between phagosome formation and nuclear lamina disassembly, we examined by live microfluidic-based confocal imaging animals expressing GFP::LMN-1, *mig-24p*::iBlueberry to mark the linker cell, as well as an mKate2-PH reporter, derived from PLC-61 and expressed in linker cell engulfing cells. mKate2-PH marks unsealed phagosome membranes by binding to phosphatidylinositol (4,5)-bisphosphate (PI(4,5)P2) [16, 23, 24]. We found that when mKate2-PH signal is present around the linker cell, revealing ongoing phagocytosis, GFP::LMN-1 remains localized to the nucleus (Figure 2A; note double membranes labelled by mKate2-PH corresponding to the engulfing cell plasma membrane and the forming phagosome membrane). Following mKate2-PH signal disappearance, signifying phagosome sealing (Figure 2B; only engulfing cell plasma membrane labelled), GFP::LMN-1 localization becomes diffuse (Figure 2C). Thus, phagosome sealing precedes linker cell nuclear lamina degradation. Of note, the time gap between mKate2-PH loss and GFP::LMN-1 redistribution is highly variable (2min, 15min, 30 min, 1h, 2h in the five animals we examined, Supplemental Figure 2), suggesting that phagosome sealing is unlikely to directly instruct nuclear lamina dismantling.

**Figure 2.**
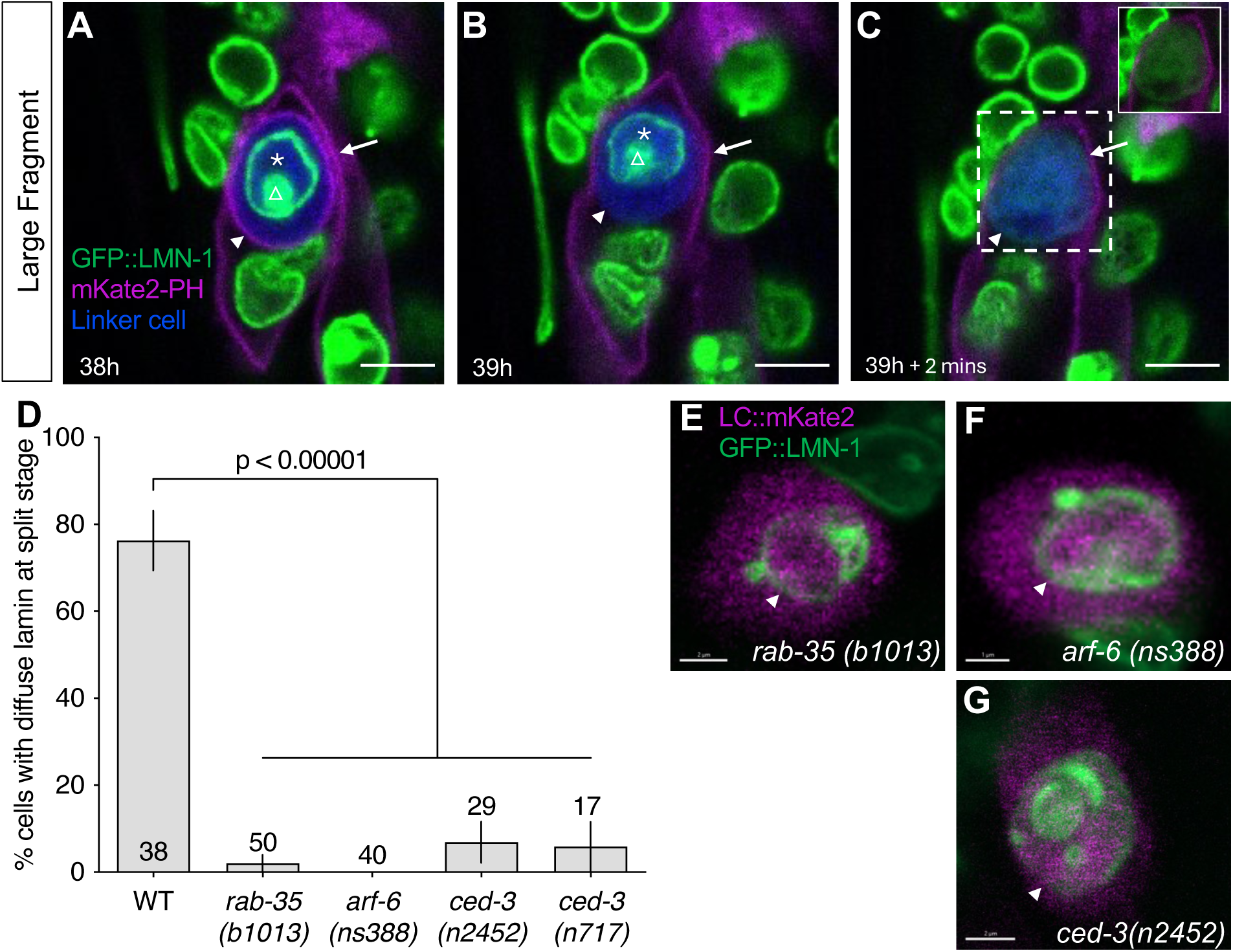
Lamina disassembly takes place upon phagosome sealing and is blocked in *rab-35*, *arf-6* and *ced-3* mutants. (A-C) Confocal live-imaging of a linker cell labeled with *mig-24*p::iBlueberry, lamin endogenously tagged with GFP, and mKate2-PH labeling unsealed phagosome membranes and plasma membrane of the engulfing cells. Asterisks, intact linker cell nuclear lamina. Arrowhead, unsealed phagosome membrane (A), unlabeled sealed phagosome membrane location (B). Arrow, engulfing cell membrane. Triangle, deformed nuclear lamina. (C) GFP alone. Scale bar, 4 μm. (D) Quantification of % linker cells with diffuse lamina at the split stage in indicated genotypes. Number of animals scored is inside/above bars. Average of independent experiments: 2 (*arf-6*); 3 (WT, *ced-3(n717)*) 4 *(rab-35)*; 5 (*ced-3(n2452)*). Error bars, standard error of sample proportion, Fisher’s exact test. (E-G) Confocal imaging of lamina status using endogenously tagged GFP::LMN-1 reporter in persisting linker cells labeled by *mig-24*p::mKate2 in 24h old adults of indicated genotypes. Scale bar, 2 μm (E,G), 1 μm (F).

To further examine the connection between phagosome formation and linker cell nucleus degradation, we examined *rab-35(b1013)* GTPase mutants in which cell splitting is unperturbed but phagosome maturation fails to initiate [16]. While 76% of wild-type animals harboring a split linker cell also exhibit GFP::LMN-1 redistribution (n=38), only 2% of *rab-35(b1013)* mutants show localization changes (Figure 2D; n=50). Nuclear localization of GFP::LMN-1 is still evident in *rab-35(b1013)* mutants observed 24 hours after linker cell death onset (10/10 animals observed; Figure 2E). We previously showed that RAB-35 inhibits the ARF-6 GTPase by promoting its removal from the phagosomal membrane [16]. We found that *arf-6(ns388gf)* GTPase mutants, which have the same linker cell clearance defect as *rab-35(b1013)* mutants [16], are also unable to redistribute GFP::LMN-1 (0%, n=40; Figure 2D), and GFP::LMN-1 remains at the nuclear membrane even 24 hours after linker cell death initiation (Figure 2F). Thus, nuclear lamina disassembly during linker cell death requires the RAB-35 phagosome maturation initiation factor acting through activation of ARF-6 [16].

### CED-3 caspase and CED-4/Apaf1 promote linker cell degradation and nuclear lamina dismantling

To identify additional regulators of linker cell nuclear degradation, we examined the effects of mutations in known linker cell death and apoptosis genes at different time points during linker cell death. Consistent with previous findings [14], linker cell death is initiated normally in *ced-3(n2452)* deletion mutants (Supplemental Figure 3A), where it is accompanied by nuclear crenellations (Supplemental Figure 3B). Unexpectedly, we found that a linker cell corpse persists in 56% (n=114) of *ced-3(n2452)* mutants even 24 hours after cell death initiation (Figure 3A). We used confocal imaging of animals carrying a *lag-2p*::GFP reporter labeling the linker cell and a *lin-48p*::mCherry reporter labeling the engulfing U.I/rp cells to score whether these persisting corpses were engulfed. In the few wild-type animals with remaining corpses, about half are engulfed (Figure 3B); however, in *ced-3(n2452)* animals, most persisting cells are not engulfed (Figure 3B,C, arrow points to the portion of the linker cell that is not engulfed; Supplemental Movie 1).

**Figure 3.**
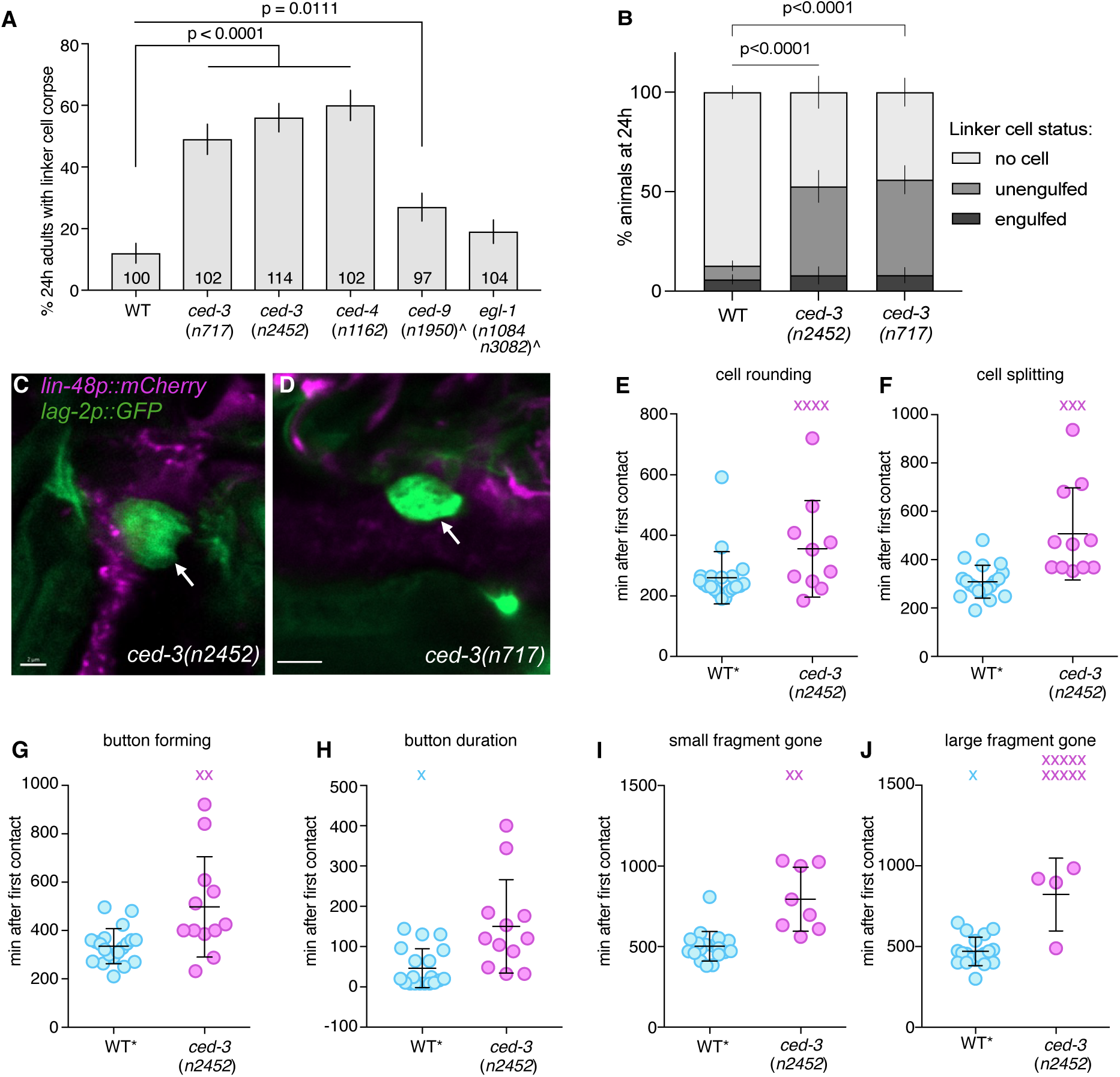
*ced-3* and *ced-4* are required for linker cell engulfment. (A) Linker cell degradation in indicated genotypes. Strains contain either *lag-2p::GFP* linker cell reporter or *nsIs1(lag-2p::GFP, rol-6(+))* (denoted by ^) and *him-5(e1490).* Number of animals scored inside bars. Error bars, standard error of sample proportion. Fisher’s exact test. (B) Linker cell engulfment in 24h adults in indicated genotypes. Strains carried *lin-48p*::mCherry engulfing cells reporter, *lag-2p*::GFP linker cell reporter. Number of animals scored: WT n=102, *ced-3 (n2452)* n=38, *ced-3(n717)* n=50. Biological replicates: 2 (WT), 3 (mutants). Error bars, standard error of sample proportion. Fisher’s exact test. (C and D) Representative confocal images of unengulfed linker cells in 24h adults in *ced-3(n2452)* and *(n717)* mutants. *lin-48p*::mCherry reporter marks engulfing cells, *lag-2p*::GFP marks linker cells. Arrow points to the unengulfed side of the linker cell. Scale bar, 2 μm (C), 3 μm (D). (E-J) Quantification of indicated parameters in long-term live-imaging of *ced-3*(*n2452*) animals. Strains contain linker cell reporter (*mig-24p::Venus*), U.l/rp cell reporter (*lin-48p::mKate2*), and *him-5*(*e1490*) to enrich for males. Dots, individual events in single animals in min with respect to first contact. X, event did not occur and was not factored into statistical analysis. Bars, mean ± std. Student’s t-test. Wild type (WT) are same quantifications as in [16].

We observed the same set of defects in *ced-3(n717)* mutants (49% persisting corpse, n=102), confirming that these are due to loss of CED-3 function (Figure 3A, B, D). Similar persisting cell defects are also observed in *ced-4*/Apaf1 mutants and to a lesser extent in *ced-9(n1950)* mutants carrying a gain-of-function mutation in the *ced-9*/Bcl2 gene (Figure 3A).

We used long-term live imaging of *ced-3(n2452)* animals, also containing a *mig-24*p::Venus linker cell reporter and a *lin-48*p::mKate2 engulfing cell reporter, to gain more insight into the temporal progression leading to the engulfment defects we observed. We found that onset of nuclear crenellation and lodging of the linker cell between the two engulfing U.l/rp cells and are the same as in wild-type animals (Supplemental Figure 3B,C). Likewise, development of the male tail, including tail tip retraction onset, appearance of first tail rays, and cuticle shedding, is normal in *ced-3(n2452)* animals (Supplemental Figure 3D-F). However, linker cell rounding and splitting are delayed (Figure 3E,F), and the formation of the refractile button-like nucleus-containing corpse is delayed as is its persistence time (Figure 3G,H). While small fragment clearance, when it occurs, is normal (Figure 3I; Supplemental Figure 3G), large fragment clearance often fails to take place (Figure 3J).

Importantly, in *ced-3(n2452)* and *ced-3(n717)* animals with persistent linker cell corpses, GFP::LMN-1 remains associated with the nuclear envelope (Figure 2D,G). This nuclear lamina degradation defect is similar to that of *rab-35(lf)* and *arf-6(gf)* mutants (Figure 2D-F). However, double mutants containing the *arf-6(gf)* mutation and either *ced-3(n2452)* or *ced-3(n717)* mutation have more severe linker cell clearance defects than either mutant alone, suggesting that *ced-3* acts, at least in part, in parallel to the ARF-6 pathway (Supplemental Figure 3H).

Together, our findings suggest that the apoptotic caspase CED-3 and its regulators, while not required for linker cell death *per se*, play auxiliary roles in linker cell degradation and nuclear lamina disruption.

### CED-3 caspase and CED-4 Apaf1 are expressed and function in the linker cell

To determine the site of CED-3 action during linker cell degradation, we examined animals carrying a *ced-3p*::*ced-3-GFP* translational reporter. We found that GFP fluorescence is not evident in the migrating linker cell or in the U.l/rp engulfing cells. However, nuclear GFP puncta and diffuse cytoplasmic fluorescence are evident in the dying linker cell (Figure 4A). Linker cell-specific expression is also seen in animals carrying *ced-3*p::GFP transcriptional reporter (Supplemental Figure 4A), consistent with the notion that CED-3 plays a cell autonomous role in linker cell clearance. Of note, GFP fluorescence is lower compared to that seen in cells dying by apoptosis in the male tail (Supplemental Figure 4A). Expression of a *ced-4p*::*mKate2* transcriptional reporter is also detected in the dying linker cell (Figure 4B). However, unlike the *ced-3* reporter transgenes we examined, *ced-4*p::*mKate2* is expressed in both the migrating and dying linker cell. Thus, *ced-3* and *ced-4* may function in the linker cell for its clearance.

**Figure 4.**
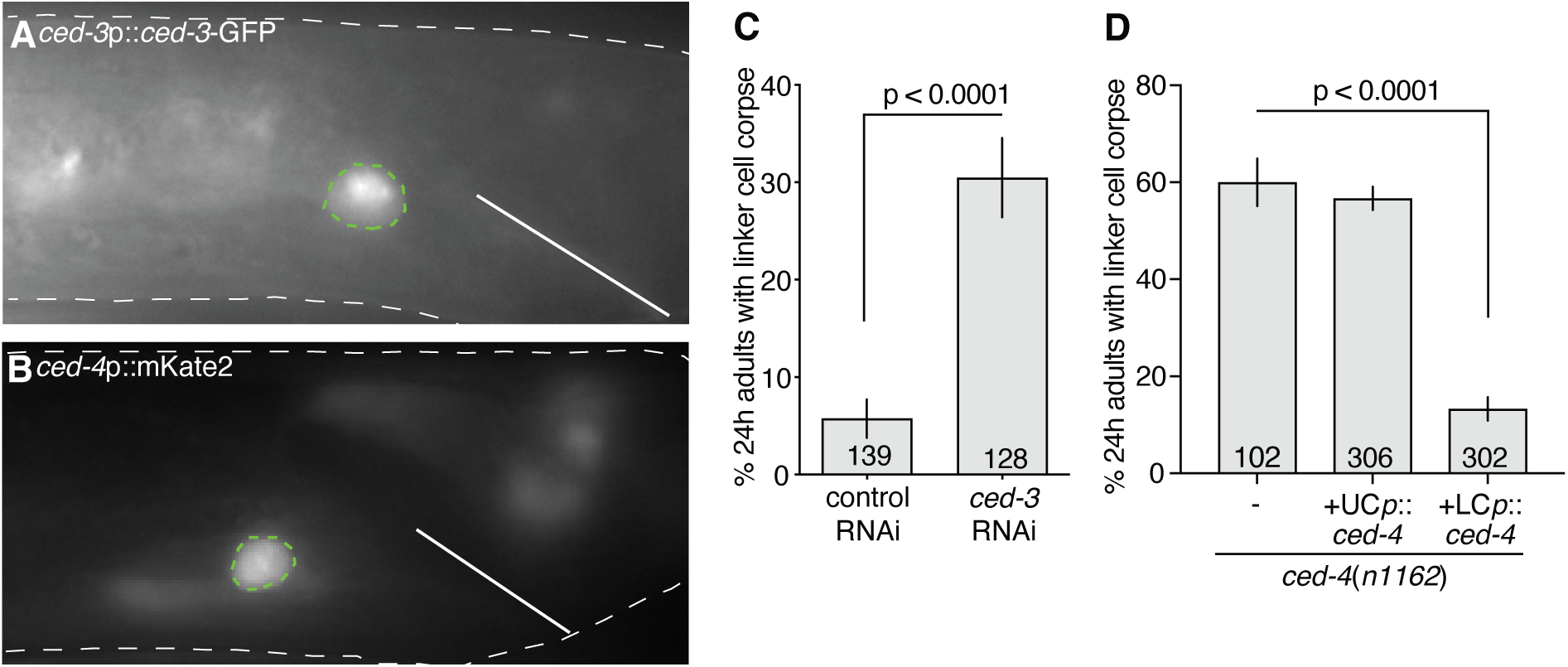
CED-3 and CED-4 function in the linker cell. (A) *ced-3* translational reporter expression and localization. Linker cell outlined in green, animal outlined in white. Solid line, position of the cloaca. (B) *ced-4* transcriptional reporter expression. Linker cell outlined in green, animal outlined in white. Solid line, position of the cloaca. (C) Persistent linker cells in 24h adults in a linker cell-specific RNAi strain. The number of animals scored is inside bars. Average of three independent experiments. Error bars, standard error of sample proportion, Fisher’s exact test. (D) Linker cell corpse persistence in indicated genotypes. Strains contain *lag-2*p*::GFP* linker cell reporter and *him-5*(*e1490*). UCp, *lin-48*p. LCp, *mig-24*p. Number of animals scored inside bars. Average of at least three independent lines. Error bars, standard error of sample proportion, Fisher’s exact test.

To further test this idea, we used linker cell-specific RNAi to inhibit CED-3 expression and found significant linker cell corpse persistence 24 hours after linker cell death initiation (Figure 4C). Furthermore, expressing *ced-4* cDNA under the control of the *mig-24p* linker cell-specific promoter rescues the linker cell clearance defect of *ced-4(n1162*) mutants; but expressing *ced-4* using the *lin-48p* engulfing cell promoter does not (Figure 4D).

Our functional studies and expression results, therefore, demonstrate that *ced-3* and *ced-4* act cell autonomously within the linker cell to promote its clearance.

### CED-3 is required for linker cell splitting and NMY-2/type II myosin shunting into the small fragment

While imaging *ced-3(n2452)* mutants with our long-term imaging setup, we noted that linker cell splitting into two fragments is significantly prolonged (Figure 3F). Furthermore, while splitting eventually proceeds to completion in some animals, in others the two fragments remain connected for hours (Figure 5A-D n=3/5 animals) with confocal imaging revealing an extended cellular bridge (Figure 5E). Consistent with these findings, imaging a synchronized population of *ced-3(n2452)* males at different timepoints reveals a significant proportion of animals with linker cell fragments connected by an unsevered bridge (“almost split” phenotype, Figure 5F). This is significantly different from a similarly synchronized wild-type male population, in which linker cell splitting is rarely observed, suggesting that it is normally a rapid process (Figure 5F; [16]).

**Figure 5.**
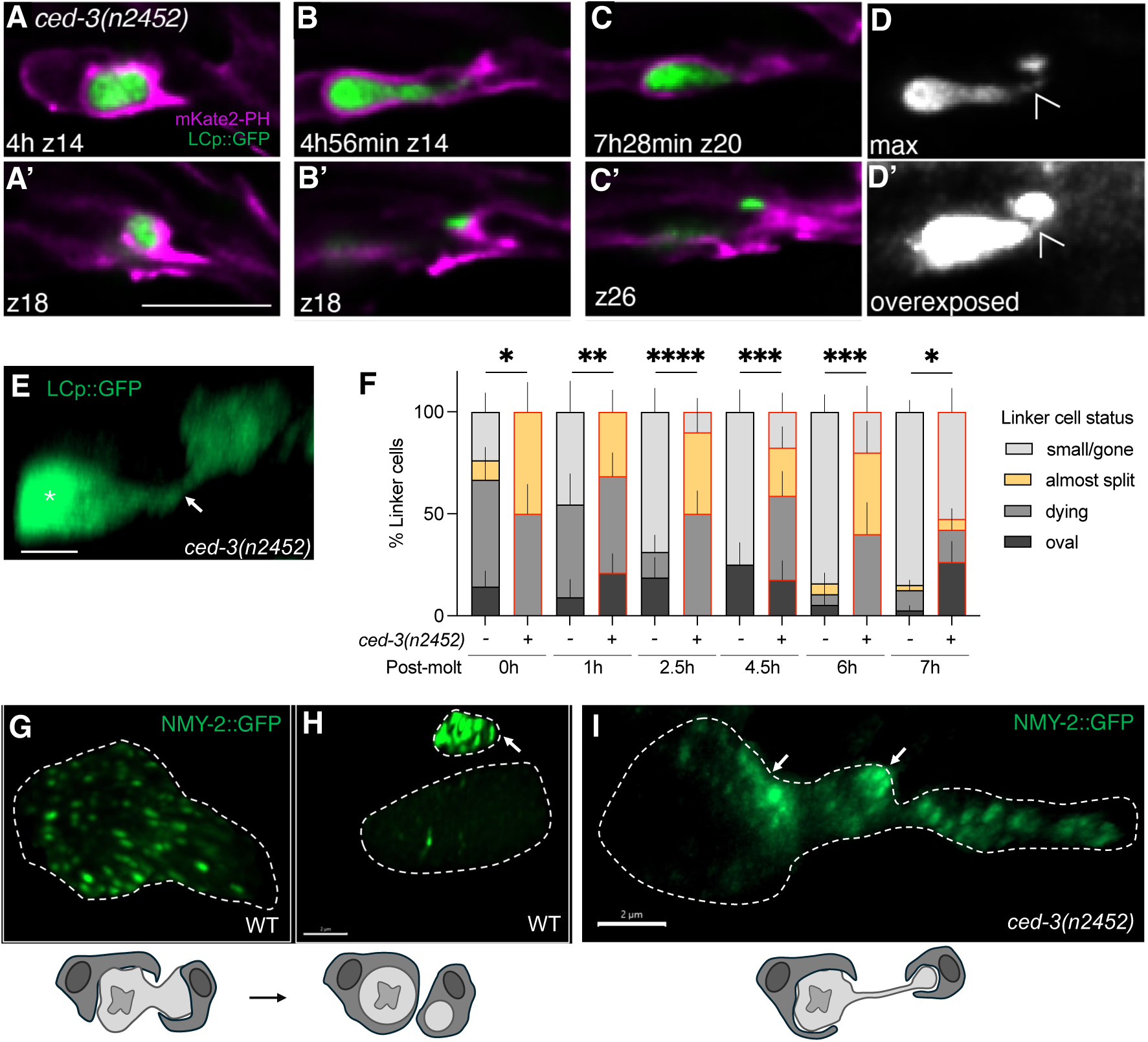
CED-3 drives linker cell splitting and NMY-2 localization. (A-D’) Long-term live imaging at two focal planes (slice number indicated by z14, z18, z26) of linker cell large fragment (top) and small fragment (bottom). Arrowhead, connecting bridge. (E) Linker cell arrested at split stage. Arrow, cytoplasmic bridge. (F) Timelines of linker cell death stages in wild type (-) and *ced-3(n2452)* (+) mutants scored by confocal imaging. Between 10 and 40 animals were scored for each timepoint. Error bars, standard error of sample proportion, Fisher’s exact test. * (p ≤ 0.05), ** (p ≤ 0.01), *** (p ≤ 0.001), and **** (p ≤ 0.0001). (G,H) Confocal imaging of NMY-2::GFP endogenous reporter showing NMY-2 localization before (G) and after (H) split stages. Scale bar, 2 μm. (I) *ced-3(n2452)* linker cell exhibits symmetric NMY-2 localization around the scission site (arrows). Scale bar, 2 μm.

Cell splitting and nuclear lamina dismantling are hallmarks of mitotic cell division, raising the possibility that linker cell fragmentation may result from an atypical cell division. To test this, we performed a candidate RNAi screen against cell cycle and other mitotic regulator genes (16 cell cycle genes, 71 gene in total). We found no effect on linker cell death initiation in any of these. However, RNAi against several candidates, including *plk-1*, *cye-1*, *nop-1* and *nmy-2*, results in cell clearance delay in animals scored 8h after the larva-to-adult molt (Supplemental Figure 5A). When examining animals expressing an endogenously tagged NMY-2::GFP, marking a type II myosin required for contractile ring formation during cytokinesis [25, 26], we noticed sequestering of GFP fluorescence to the small fragment during linker cell splitting (Figure 5G,H, n=16). By contrast, in *ced-3*(*n2452*) animals, NMY-2::GFP fluorescence is evident in both fragments and localizes near and within the connecting cellular bridge (Figure 5I, n=9/13). During mitosis, microtubules assemble into spindles. Interestingly, we also observed microtubule reorganization during LCD, visualized by a tubulin TBA-1::TagRFP reporter; however, instead of forming spindles, microtubules form condensed puncta (Supplemental Figure 5B-E’).

We conclude, therefore, that CED-3 caspase is required for linker cell splitting and NMY-2 redistribution, and that failure of these events may account for the subsequent defects in linker cell engulfment and nuclear lamina degradation. Our results also raise the possibility that linker cell death may be accompanied by an atypical form of cell division.

### Mouse embryonic fibroblasts treated with staurosporine undergo non-apoptotic cell death reminiscent of linker cell death

Staurosporine (STS), a non-specific kinase inhibitor [27], has been suggested to induce apoptosis in cultured cells [28, 29]. However, staurosporine-treated immortalized fetal thymocytes from mice lacking *Apaf1*, *Casp9*, *Casp8*, or *Bax* and *Bak* still die, and death is accompanied by caspase 3 processing [21], suggesting possible similarities with linker cell death. To expand on the thymocyte studies, we treated wild-type or *Casp3; Casp7^DKO^* mouse embryonic fibroblasts (MEFs) with STS or DMSO control. While both caspases are activated in the wild type, as monitored by caspase auto-cleavage and PARP substrate cleavage (Supplemental Figure 6A,B), death proceeds even in the absence of Caspase 3 and 7 (Figure 6A). Although caspase-deficient cells die more slowly, by 77h of treatment, overall cell death is comparable to wild-type cells (Figure 6A). Thus, STS-induced death has a Caspase-3/7 independent component. Strikingly, STS treatment of either wild-type or *Casp3; Casp7^DKO^* MEFs results in nuclear crenellations observed by lamin B1 immunofluorescence (Figure 6B-E) and by electron microscopy (Figure 6F-I). Bulk RNAseq on wild-type fibroblasts at 30 min following STS exposure, to uncover early acting genes, reveals induction of a ubiquitin proteasome system component (*Tdpoz8*) as well as of members of the ALYREF mRNA export complex (Figure 6J, Supplemental Figure 6C,D, Supplemental Table 1).

**Figure 6.**
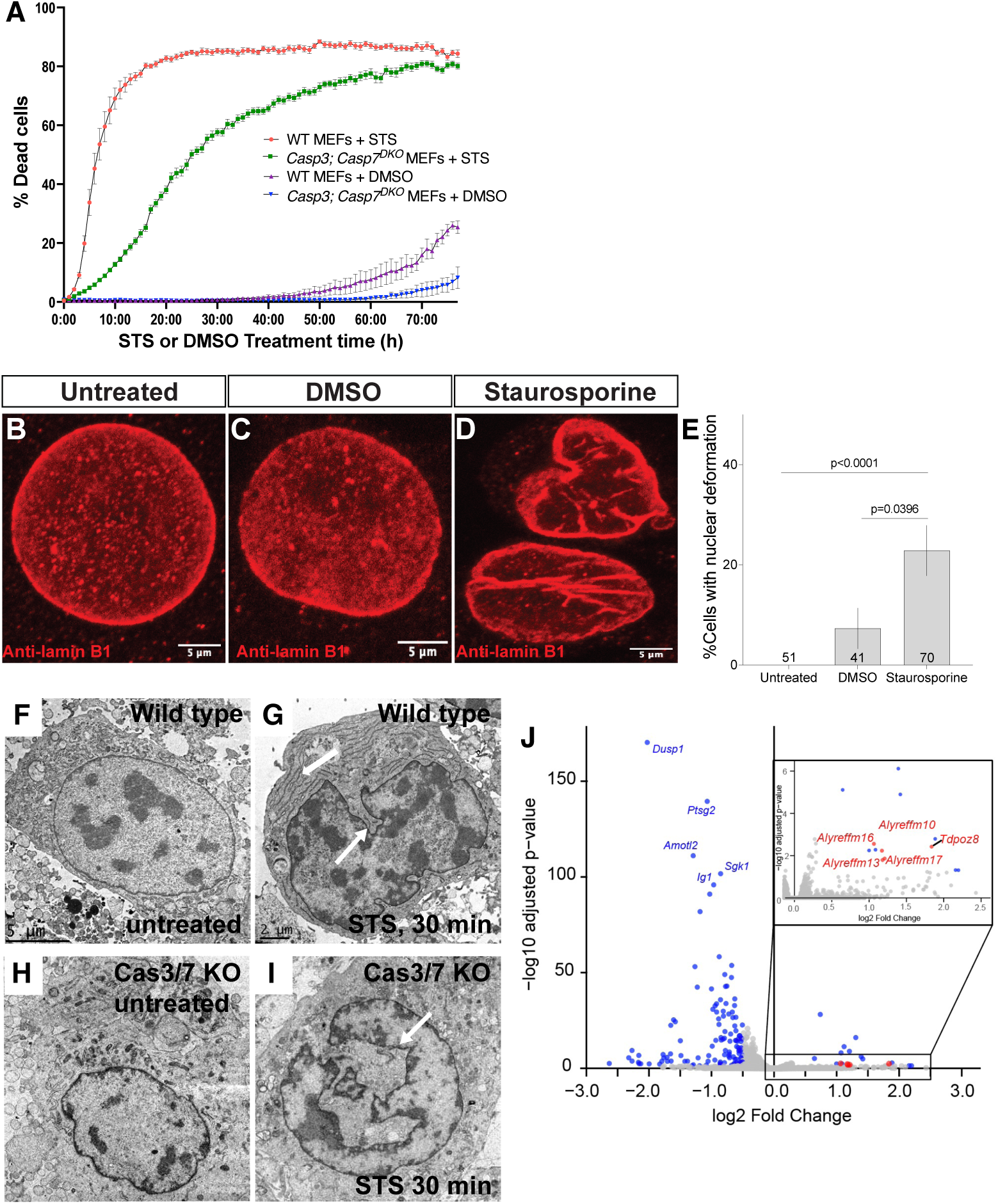
Mouse embryonic fibroblasts (MEFs) acquire LCD hallmark nuclear deformations accompanied by cell death in a caspase-independent manner upon treatment with staurosporine (STS). (A) Caspase 3/7 KO MEFs treated with staurosporine (STS) undergo caspase-independent cell death. Average of three independent experiments. (B-D) Representative immunofluorescence images of lamin B1 staining of wild-type (WT) MEFs before treatment (A) and after treatment with either DMSO (B) or 1.5 µM staurosporine (C) for 2 h. (E) Quantification of cells positive for nuclear deformation, shown as a percentage. Number of animals scored inside bars. Error bars, standard error of sample proportion, Fisher’s exact test. (F-I) Ultrastructural details of wild-type and apoptotic mutant MEFs treated with STS. Arrows denote nuclear deformations. (J) Volcano plot displaying significantly differentially expressed genes; padj < 0.05 & log2FoldChange < +/-0.5).

Together, these findings uncover morphological similarities between linker cell death and STS-induced death of MEFs, and raise the possibility that similar mechanisms may drive cell demise in these settings.

## Discussion

### CED-3 may have a primary role in linker cell splitting

The findings described in this study are consistent with a model in which non-apoptotic death of the *C. elegans* linker cell requires components of the apoptotic machinery, including the principal caspase, CED-3, to execute auxiliary tasks that contribute to the fidelity of dying cell degradation. Indeed, loss of CED-3 caspase leads to defects in linker cell splitting, rounding -- indicative of abnormally maintained adhesion to neighboring cells, corpse engulfment, and nuclear degradation. Do these defects reflect independent functions of CED-3 caspase, or might they be related?

The earliest linker cell degradation defect we observe in *ced-3* mutants is the inability to efficiently split the cell into a large and a small fragment (Figure 7). This defect correlates with abnormal distribution of non-muscle myosin NMY-2, which instead of concentrating primarily in the small fragment, aberrantly localizes to both fragments near the presumptive scission site. Our findings support the notion that all other defects we observe in *ced-3* mutants could stem from this initial cell splitting abnormality. As we previously demonstrated, linker cell splitting occurs concomitantly with engulfment of each linker cell fragment by a different engulfing cell, either U.lp or U.rp, through competitive phagocytosis [16]. We posit that a defect in linker cell fragment separation, leading to persistence of a fragment-joining bridge, forms a physical barrier that prevents engulfing cells from fully surrounding each linker cell fragment (Figure 7). This, in turn, prevents formation of a sealed phagosome. As we show, phagosome sealing is likely a pre-requisite for nuclear dismantling, as mutants that prevent early stages of phagosome maturation fail to dismantle the linker cell nucleus. Thus, the persistence of an intact nucleus in *ced-3* mutants is likely an indirect consequence of a defect in linker cell splitting (Figure 7).

**Figure 7.**
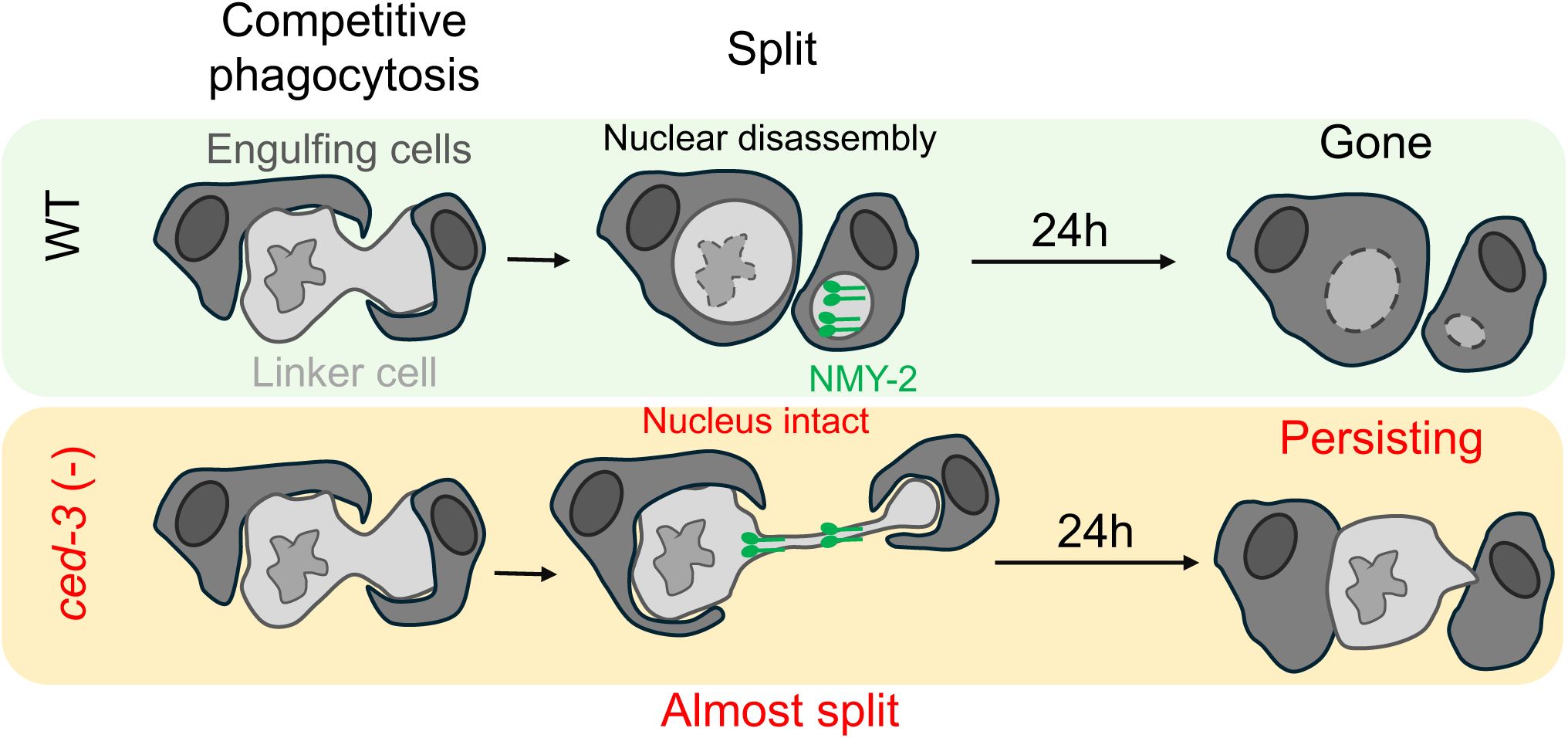
Model for CED-3 function in linker cell removal. In wild-type animals, linker cells undergo splitting, followed by lamina disassembly and asymmetric NMY-2 redistribution to the small anuclear fragment. The linker cell is then degraded. In *ced-3(-)* animals, splitting is stalled, NMY-2 localizes to both fragments and to the bridge between the two fragments. This results in failure of engulfment and nuclear dismantling.

### Is linker cell death a consequence of a failure in cell division?

Caspases carry out numerous non-apoptotic tasks. In *C. elegans*, for example, CED-3 controls p38 MAPK activity during larval development [30]. Caspases also ensure the fidelity of cell division. Mammalian Caspase 2, for example, is important for cell cycle progression in mouse embryonic fibroblasts [31], and Caspase 4 promotes epithelial cell division by controlling actin polymerization [32]. Other caspases have also been suggested to control mitosis [33]. These observations raise the intriguing possibility that linker cell splitting, which requires CED-3 caspase activity, is mechanistically related to cell division. Our other findings are consistent with this idea. Nuclear lamina degradation, for example, is a common feature of both mitosis and linker cell death [34]. In addition, localization of non-muscle myosin to the cytokinetic furrow in mitotically dividing cells and to the linker cell scission site may reflect similar functions [35]. Reorganizing of tubulin also occurs during mitosis and, in a different way, in linker cell death. Furthermore, as in many other non-apoptotic roles for caspases, the level of caspase expression in non-apoptotic settings, including cell division, is an order of magnitude lower than in dying cells [36-38]. The same appears to be the case, as we describe here, in the linker cell.

The juxtaposition of an apoptotic caspase, CED-3, with a non-apoptotic role in cell scission, during a non-apoptotic cell death process, LCD, is intriguing and may point to the evolutionary origins of LCD and apoptosis. In one model, LCD may be an ancient cell death program activated by a failure in cell division, a process that was perhaps once primarily governed by caspases. As redundant cell division mechanisms arose, caspases were instead co-opted to play a more central role in cell death, leading to the development of apoptosis. Caspase-dependent apoptotic cell death appears to be a much more efficient process than LCD [2], supporting an evolutionary incentive to favor a role for these cysteine proteases in cell death.

### Is the linker cell-type death mechanism conserved in mammals?

We previously published a survey of electron micrographs demonstrating that cell death accompanied by nuclear crenellations is commonplace in vertebrate development and in disease [3]. Our finding that MEFs undergo a similar morphological change in response to treatment with the cell-death inducer and kinase inhibitor staurosporine adds to this notion. Although our findings suggest involvement of a ubiquitin proteasome system factor early on in staurosporine-induced death, whether this factor is functionally required for LCD remains to be determined. Nonetheless, the induction of cleaved CASP3 and PARP following staurosporine treatment, despite the ability of cell death to proceed in *Casp3/7* double knockout cells, suggests molecular similarities with linker cell death. Disruption of these and other genes identified in our RNAseq studies may reveal the extent of molecular conservation with *C. elegans* linker cell death.

## Acknowledgements

We thank Cori Bargmann and Shaham lab members for comments on the manuscript. We would like to acknowledge the Rockefeller University’s Bio-Imaging Resource Center, RRID:SCR_017791 and the Electron Microscope Imaging Facility of CUNY Advanced Science Research Center for instrument use and scientific and technical assistance. We thank Dr. B. Shklyar and L. Simchi from the Bioimaging Unit at the University of Haifa for their guidance and assistance. We thank Verena Jantsch-Plunger for strains and Orna Cohen-Fix for discussion. Some strains were provided by the CGC, which is funded by the NIH Office of Research Infrastructure Programs (P40 OD010440). This work was supported by NIH NRSA Training Grant GM066699 to L.M.K., NIH K99GM151467 to O.Y., and NIH grants R35NS105094 and R01HD103610 to S.S.

## Author contributions

Conceptualization, O.Y, L.M.K. and S.S.; Methodology, O.Y, L.M.K., W.K., B.O.B., J.K., D.M., Y.L., S.L. and S.S.; Software, W.K.; Formal Analysis, O.Y., L.M.K., J.K., Y.K., Y.L., E.P.; Investigation, O.Y, L.M.K., B.O.B., J.K., D.M., Y.K., Y.L.; Writing – Original Draft, O.Y., L.M.K., S.S.; Writing – Review & Editing, O.Y., L.M.K., W.K., S.L. and S.S.; Funding Acquisition, O.Y., L.M.K., W.K., and S.S.; Resources, O.Y., L.M.K., W.K., B.O.B., J.K., S.L., and S.S.; Visualization, O.Y., L.M.K., W.K.,Y.L., E.P., Y.K., J.K., and S.S.; Supervision, S.L. and S.S.

## Materials and Methods

### Key reagent table

Key reagent table.
| Resource | Source |
| --- | --- |
| *nsIs589*: UV/TMP integration of *nsEx5123* (pLMK09[*lin-48p::mKate2*][7] (10 ng/ul), *unc-119(+)* 25 ng/ul, pBluescript (65 ng/ul)] | [22] |
| *bcls109*: *ced-3p::ced-3-GFP*, rol-6(+), | gift from B. Conradt, [39] |
| *nsEx5307*: pLMK65(*ced-4p::mKate2*) (25 ng/ul) + *unc-119(+)* (25 ng/ul) + pBluescript (50 ng/ul)] | This study |
| *nsEx5627,-8,-9*: pLMK66(*mig-24p::ced-4a* cDNA) (20 ng/ul) + *odr-1p::RFP(+)* (20 ng/ul) + *pEB30* (20 ng/ul) + pBluescript (40 ng/ul) | This study |
| *nsEx5619,-20,-81*: pLMK67(*lin-48p::ced-4a* cDNA-SL2-mCh) (20 ng/ul) + *odr-* | This study |
| *1p::RFP(+)* (20 ng/ul) + pBluescript (60 ng/ul) |  |
| *nsIs495*: UV/TMP integration of *nsEx4360(lin-48p::mCherry* (25 ng/ul)+ pBluescript (75 ng/ul)) | This study |
| *nsIs25*: *ced-3p::GFP, rol-6(+)* | gift from C. Maurer, [40] |
| *nsEx4944*: *pPG08(ced-3p::mCherry)* (10 ng/ul) + *unc-119(+)* (25 ng/ul) + pBluescript (65 ng/ul) | gift from P. Ghose |
| *nsIs653*: UV/TMP integration of *nsEx5304(lin-48p::mKate2-PH* (25 ng/ul)+ pBluescript (75 ng/ul)) | [22] |

### *C. elegans* husbandry and strains

*C. elegans* strains were raised at 20°C on nematode growth medium (NGM) seeded with OP50 bacteria, unless otherwise indicated. Wild-type animals were of the Bristol N2 strain. All strains carry a *him-5*(*e1490*) mutation that generates a high incidence of male progeny. Transgenic lines were generated by injection of plasmid DNA mixes into the hermaphrodite gonad. Integrated transgenic strains were generated with UV/trioxalen treatment (Sigma, T6137) [41] and were outcrossed at least four times before scoring. Most strains also have one of four integrated linker cell markers: *nsIs763*(*mig-24p::*iBlueberry) II*, qIs56*(*lag-2*p::GFP) V, *nsIs65(*[*mig-24*p::Venus) X, or *nsIs650(*[*mig-24*p::mKate2) X. Other alleles are as follows:

LGI

*lmn-1(jf98), nmy-2(cp13), tba-1(pg77)*

LGII

bqSi142 (*emr-1p*::*emr-1*::mCherry + *unc-119*(+));

LGIII

*ced-9(n1950), ced-4(n1162), rab-35*(*b1013*), *rab-35*(*tm2058*), *tbc-10*(*gk388086*), *unc-119*(*ed3*)

LGIV

*ced-3(n717), ced-3(n2452), arf-6*(*tm1447*), *arf-6*(*ns388*)

LGV

*egl-1*(*n1084n3082*)

LGX

*stIS10116*(*his-72p*::*his-24*::mCherry::*let-858* 3’UTR)

### Germline Transformation and Rescue Experiments

Plasmid mixes containing the plasmid of interest, co-injection markers, and pBluescript were injected into both gonads of young adult hermaphrodites [42]. Injected animals were singled onto NGM plates and allowed to grow for two generations. Transformed animals based on co-injection markers were picked onto single plates, and screened for stable inheritance of the extrachromosomal array. Only lines from different P0 injected hermaphrodites were considered independent. For cell-specific rescue experiments, animals expressing fluorescence protein in either the linker cell or the U.l/rp cells were picked to a new plate at the early L4 stage, before linker cell death, to avoid bias. Isolated animals were then staged based on tail morphology under DIC microscopy for appropriate linker cell scoring.

### Linker Cell Survival and Corpse Persistence Assays

Linker cell death was scored as previously described [19]. Briefly, unstarved gravid hermaphrodites were bleached to isolate embryos, which were allowed to hatch overnight in M9. Synchronized L1 animals were released on 9-cm NGM plates seeded with OP50 or HT115 *E. coli* containing the RNAi construct of interest on IPTG-RNAi plates. Male animals were isolated onto a new plate prior to the L4-to-adult transition based on full retraction of the male tail tip with rays visible under the unshed L4 cuticle. 2h later (linker cell survival) or 8h and 24h later (corpse persistence), these animals were mounted onto 2% agarose-M9 pads, anaesthetized in 25 mM sodium azide, and examined on an Axioplan 2 fluorescence microscope (Zeiss) with a 63x/1.4NA objective (Zeiss) and DIC optics. The linker cell was identified by location, morphology, and fluorescence from reporter transgenes. Cells were then scored as surviving (normal nucleus), dead (deformed nucleus, competitive phagocytosis, split or rounded stages), or gone based on DIC morphology and fluorescence.

### Imaging LCD stages

Animals were synchronized by bleaching as described above. Appropriate timecourse was determined for each genotype/strain. Generally, animals were images on a second day after plating (36-46h post-plating/post L1 release). LCD stage was determined by the shape of the linker cell: oval – oblong, not rounded shape; competitive phagocytosis – budding shape; split – large and small fragments visible; round – refractile corpse; almost split – two fragments connected by a long cytoplasmic bridge (Figures 1, 5, 7). “Almost split” phenotype was carefully scored by confocal imaging only.

### Engulfment scoring

Engulfment of the linker cell (Figure 3B-D) was scored by confocal microscopy using Zeiss LSM 900 inverted laser scanning confocal microscope with a 63x/1.4 NA oil objective. Linker cell was labeled with *lag2p*::GFP, while engulfing cells were labeled with *lin-48p*::mCherry. Linker cell was considered engulfed if in all the z stacks it was surrounded by engulfing cells. Linker cell was scored as unengulfed if it had exposed/engulfing cell-free side in any of the z stacks. Animals were scored in 24h synchronized adults.

### Linker cell-specific RNAi

RNAi was performed by feeding on freshly made (within two weeks) plates containing 1mM IPTG and 25μg/ml cabenicillin [18] using a linker cell-specific RNAi strain (OS7637, genotype: *him-8*; *rde-1(ne219)* qIS56=*lag2p*::GFP (V); nsIs387=mig-24p::rde-1::SLC-mCherry + lag-2::mCherry. Gravid hermaphrodites were bleached and the released embryos were synchronized at the L1 stage by leaving them overnight in M9. L1 animals (30% of which are male) were added to each RNAi plate and grown for approximately 48h at 20°C. Animals were picked at the L4-adult molting stage and 2h or 8 hr adults were mounted onto 2% agarose-M9 pads and immobilized using 25mM sodium azide. Linker cell presence and morphology was scored using an upright fluorescent microscope Zeiss Axio Scope.A1 with a 63x/1.4NA objective and DIC optics. Reagents were checked by performing GFP RNAi experiment, and GFP fluorescence knock-down was scored in late migrating linker cells (n=10). At least 60% knockdown (6 out of 10 animals) was considered a good positive control. Empty vector control was used as a negative control, and was always scored on the same day with other gene knock-downs to validate the baseline. Clones were from the Ahringer feeding library [43] confirmed by Sanger sequencing.

### Electron Microscopy

2h adult males with a linker cell GFP marker were mounted on 2% agarose/M9 pads and lightly anesthetized with 25mM sodium azide, then observed under a fluorescence dissecting microscope or by DIC microscopy to determine the approximate stage of linker cell death. Split stage accompanied by lamina dissociation was verified by the presence of perinuclear blebbing (Supplemental Figure 1C). Animals were then fixed, stained, embedded in resin, and serially sectioned using standard methods [44].

MEFs were grown on grids in standard media, fixed, stained, embedded and serially sectioned for electron microscopy using standard methods [44]. Serial images were acquired using a Titan Themis 200 kV transmission electron microscope with Cs Image Corrector. Image processing and analyses were performed using ImageJ and IMOD software.

### Confocal Imaging

Confocal images were acquired using a Zeiss LSM 900 inverted laser scanning confocal microscope with a 63x/1.4 NA oil objective. Animals were immobilized in 200 mM sodium azide either on 1% agar pads or using microfluidic devices [22].

### Long-Term Imaging and Movie Generation

Early L4 male animals were imaged in a microfluidic device [22] for Figures 1F, 2A-C, 3E-J, S2, S3B-G, and 5A-D’. Animals were fed a constant flow of NA22 bacteria in S medium, supplemented with kanamycin (50 ng/μl) to prevent bacterial overgrowth. Animals were immobilized and imaged every 8 minutes for at least 20h. Mutant animals were typically imaged for >30h. Exposure time and light intensity was held constant across strains when the same integrated transgene was imaged. Occasionally tail development was perturbed by the flow of medium and repeated immobilization procedures, and these animals were not included in subsequent analyses.

Confocal long-term imaging was performed manually (Figures 1F, 2A-C) over 10-12h. Animals were immobilized every 30 minutes to check the stage of the linker cell. Images were taken more frequently around the stages of interest. Exposure time and light intensity were held constant.

### Deconvolution

Deconvolution was performed using Hyugens Pro with theoretically calculated PSFs, all auto options, except the number of iterations was increased to 200.

### Image Analysis

For long-term imaging: Image analysis was performed using custom-written MATLAB R2016b (Mathworks) scripts [22]. To overlay imaging frames, we straightened each three-dimensional image stack using a previously published algorithm [22, 45] based on a manually selected worm backbone in the DIC channel. To correct for small residual animal movements during multi-channel acquisition, obvious landmarks visible in the fluorescence channels, such as the cloaca, fluorescently labeled cells in the animal’s tail, or vesicles within the U-cell descendants were then manually aligned to the DIC channel in each frame. Time-lapse movies were generated by centering all straightened, aligned images on the linker cell and cropping the entire movie to an appropriate size. Frames were removed from the movies whenever residual animal movement excessively blurred the images.

For standard confocal imaging: Images were processed using Imaris version 10.12.0 or ImageJ.

### Plasmid Construction

Plasmids containing *lin-48*p, *mig-24*p, or transcriptional reporters were cloned using multi-piece one-step cloning into a modified pPD95.75 backbone lacking GFP [16].

### Quantification and Statistical Analysis

Statistical analysis was performed using GraphPad Prism. Statistical parameters including mean ± standard error of the proportion, mean ± SEM, and n are reported in the main text, figures and figure legends. Number of independent experiments are reported in the figure legends. Independent experiments are biological replicates meaning that animals were processed from different cultures on different days. Data is judged to be statistically significant when p < 0.05 by Fisher’s exact test or student’s t-test (both two-sided), where appropriate. We used sample size calculator to identify the sample of size of 30 that allows detection of meaningful difference with a power of 0.8.

Randomization: for all scoring controls and mutant strains were processed on the same day.

### Genetic Data Analysis

When comparing a transgenic line to the parental strain, a minimum of three independent lines were scored. Approximately 100 animals from each of these lines were examined for linker cell defects, and compared to the parental strain using a Fisher’s exact test to determine significance. For ease of presentation, we pooled the lines from a given experiment and displayed them on the relevant figure using mean ± SEM (N = 3, 4, or 5 lines).

### Cell Culture and reagents

MEFs WT cell were grown in complete DMEM medium (1% sodium pyruvate, 1% L-glutamate, 1% non-essential amino acids, 1% Pen-Strep, 10% fetal bovine/calf serum, and 0.1% β-mercaptoethanol [46] at 37°C. Staurosporine (STS, 1.5 µM) was purchased from Fermentek (cat#62996-74-1.5).

#### Cell line identity

Mouse embryonic fibroblasts (MEFs) from the laboratory of Dr. Larish were authenticated by genotyping and routinely tested for mycoplasma contamination every two months using a PCR-based detection assay. These cells are not listed in the ICLAC database of commonly misidentified cell lines.

### Bulk RNA-sequencing and analysis

Wild-type MEF cells were harvested 30 min after treatment with STS or DMSO. Bulk RNA was isolated with RNeasy Kit (Qiagen) following supplier’s instructions. For measurement of RNA quantity, the Agilent 2100 Bioanalyzer was used. RNA-seq libraries were processed and sequenced on a NextSeq High Output (SR 75) at the Rockefeller University Genomics Core Facility.

Sequencing was quality-checked using FastQC (v0.12.1). Reads were aligned to the *Mus musculus* reference genome (GRCm39) using Rsubread (v2.22.1), and gene-level read counts were obtained with *featureCounts*. Differential expression analysis was performed using DESeq2 (v1.48.1) in R (v4.5.1). Genes with an adjusted *p*-value (padj) < 0.05 were considered significantly differentially expressed. Genes with a log₂ fold change (log₂FC) < -1 were classified as downregulated, and those with log₂FC > 1 as upregulated. Data analysis and visualization were conducted using the tidyverse (v2.0.0) framework, plots were generated via ggplot2 (v4.0.0).

### Quantification of cell death in MEFs culture

Apoptosis was induced with 1.5 µM Staurosporine (STS) in Caspase-3/7 wild-type (WT) and Caspase-3/7 knockout (KO) MEFs. Cells treated with DMSO served as controls. Plates were immediately placed in the Incucyte Live-Cell Analysis System, and cell death progression was monitored every hour for 77 hours. Data were analyzed using the AI Cell Health Analysis tool, which performs label-free classification of live and dead cells based on machine-learning analysis of phase-contrast images. The algorithm identifies characteristic morphological features associated with cell death, including cell rounding, detachment (in adherent cells), and changes in texture and brightness. This approach enables automated and objective assessment of cell viability without reliance on fluorescent dyes. Live/Dead Classification: A separate AI model, trained on paired phase and fluorescence images of known live and dead cells, learns to differentiate dead cells based on their unique morphology (e.g., shrinkage, detachment, loss of refringence). Statistical analysis was performed using GraphPad Prism, with four to five technical replicates per condition and three independent biological experiments.

### SDS-PAGE and Western blot Analysis (Supplemental Figure 6A,B)

Cells were lysed in Whole-Cell extract buffer (25 mM HEPES, pH 7.7, 0.3 M NaCl, 1.5 mM MgCl2, 0.2 mM EDTA, 0.1% Triton X-100, protease inhibitor cocktail (Roche, 1:25 dilution), 100 μg/ml phenylmethylsulfonyl fluoride (PMSF) and 50ug/ml Dithiothreitol (DTT)) and placed on ice for 30 min (vortexing once after 15 min). After 30 min, the samples were centrifuged at 13,000 . g for 10 min at 4 °C. The supernatants containing total protein were analyzed for protein concentration using the Bio-Rad Protein Assay Dye Reagent Concentrate Kit. Proteins (50 μg) were separated by sodium dodecyl sulfate–polyacrylamide gel electrophoresis (12%SDS-PAGE), followed by transfer to a nitrocellulose membrane. Membranes were blocked with 5% (w/v) non-fat dried skimmed milk powder in PBS supplemented with 0.05% Tween-20 (PBS-T) for 1 h at RT. Next, primary antibodies were added at 4 °C overnight or for 2 h at room temperature. Membranes were then washed three times with PBS-T buffer, incubated with the secondary antibody for 1 h at RT and washed three times for 15 min each with PBS-T. Western Bright ECL (Advansta) was added to the membrane for 30–60 s and analyzed using the Image Quant LAS-4000 (GE Healthcare Life Sciences) and Image Quant LAS-4000 software (GE Healthcare Life Sciences). Densitometry of proteins levels were determined by Image studio lite v5.2. Primary antibodies: Cleaved PARP (cPARP)—rabbit polyclonal anti-PARP antibody (Cell Signaling cat#5244) at a dilution of 1:1000 for overnight. Cleaved Caspase-3—rabbit monoclonal antibody anti-active Caspase-3 Asp175 (Cell Signaling cat#cs9664) at a dilution of 1:1000 for 2h. Actin—mouse monoclonal anti-actin antibody (ImmunoTM cat#08691002) at a dilution of 1:50,000 for 2h.

### Immunofluorescence

Cells grown on 0.1% gelatin coated coverslips were fixed with 4% paraformaldehyde for 20 mins at room temperature and washed three times with DPBS for 10 mins per wash. Cells were permeabilized with 0.2% Triton X-100 in 10% normal goat serum for 15 mins, followed by three washes with DPBS for 5 mins per wash. Cells were then blocked in 10% normal goat serum for 2 hr at room temperature and incubated overnight at 4 °C with a primary antibody against lamin B1 (rabbit anti–lamin B1, 1:500; Abcam, ab16048). The following day, coverslips were washed three times with DPBS for 5 mins per wash and incubated with an Alexa Fluor–conjugated secondary antibody (anti-rabbit, Invitrogen, A-11012; 1:1000) for 1 hr at room temperature. Cells were subsequently washed three times with 1× PBST (0.05% Tween-20 in DPBS) for 5 mins per wash and incubated with Hoechst 33342 at a 1:10,000 dilution for 5 mins at room temperature, followed by three additional washes with DPBS for 5 mins per wash. Coverslips were mounted using SlowFade Gold Antifade Mountant (Invitrogen, S36940). Images were acquired using a Zeiss LSM900 scanning confocal microscope. Nuclear deformation analysis was performed using the Cell Counter plugin in Fiji software. Nuclei were classified as deformation-positive if they exhibited a line traversing the nucleus or pronounced nuclear invaginations. The percentage of deformation-positive nuclei was calculated relative to the total number of cells counted.

### SDS-PAGE and Western Blot Analysis (Supplemental Figure 6E)

Cells were lysed in Pierce RIPA Buffer (Thermo Scientific, 89900) supplemented with 0.2 mM EDTA and 100× Halt Protease Inhibitor (Thermo Scientific, 78429) on ice. After 5 mins, samples were centrifuged at 14,000 × g for 15 mins at 4 °C. The resulting supernatants containing total protein were collected, and protein concentrations were determined using the Pierce BCA Protein Assay Kit (Thermo Scientific, 23225). Proteins (40 µg) were separated by SDS–PAGE using 4–20% Tris–glycine gels and transferred overnight at 4 °C onto 0.22 µm PVDF membranes. Membranes were blocked with 5% (w/v) non-fat dried skim milk in TBS supplemented with 0.1% Tween-20 (TBS-T) for 1 hr at room temperature. Primary antibodies were then incubated with the membranes either overnight at 4 °C or for 2 hr at room temperature. Membranes were washed three times with TBS-T for 5 mins per wash, incubated with secondary antibodies for 1 hr at room temperature, and washed an additional three times with TBS-T for 5 mins per wash. Membranes were incubated with Immobilon Classico Western HRP Substrate (Millipore, WBLUC0100) for 60 secs and imaged using the Bio-Rad ChemiDoc Imaging System.

Analysis was performed using Image Lab software (Bio-Rad). Primary antibodies used were caspase-3 (anti-rabbit; Cell Signaling Technology, 9662), caspase-7 (anti-rabbit; Cell Signaling Technology, 9492), and β-actin (anti-mouse; Cell Signaling Technology, 3700). Secondary antibodies included anti-mouse IgG, HRP-linked antibody (Cell Signaling Technology, 7076) and anti-rabbit IgG, HRP-linked antibody (Cell Signaling Technology, 7074). All primary antibodies were diluted 1:1000, and secondary antibodies were diluted 1:2000.

### Antibody validation

Antibody specificity was validated in the context of each experimental application used in this study. For immunoblotting, antibodies were validated by loss of the target protein signal in knockout samples. For immunofluorescence, antibodies were used to assess nuclear envelope morphology and showed staining patterns consistent with known nuclear envelope localization.

## Data and Software Availability

Custom image analysis scripts are available upon request. FASTQ and count files for bulk RNA-seq are publicly accessible on Zenodo (10.5281/zenodo.17462830). Protocols and information about reagents are available upon request.

Randomization is not applicable. Scoring was not blinded.

## Notes

### Competing Interest Statement

The authors have declared no competing interest.

### Summary of Updates

Updated author list Updated Figure 5 Updated Figure 6 with new data

