## Supplemental Figures for "*C. elegans* CED-3 caspase promotes non-apoptotic linker cell dismantling but not death onset"

**Sup Table 1. Differentially expressed genes.** p-adj <0.05; LFC +/- 0.5

**Supp Movie 1. *ced-3* Linker Cell Death and Degradation,** Strain contains a linker cell reporter pseudocolored green (*mig-24p*::Venus), a U.l/rp cell reporter pseudocolored magenta (*lin-48*p::mKate2), and *him-5*(*e1490*). Time, hr:min after the first contact. max Z projection.

### SUPPLEMENTAL FIGURES

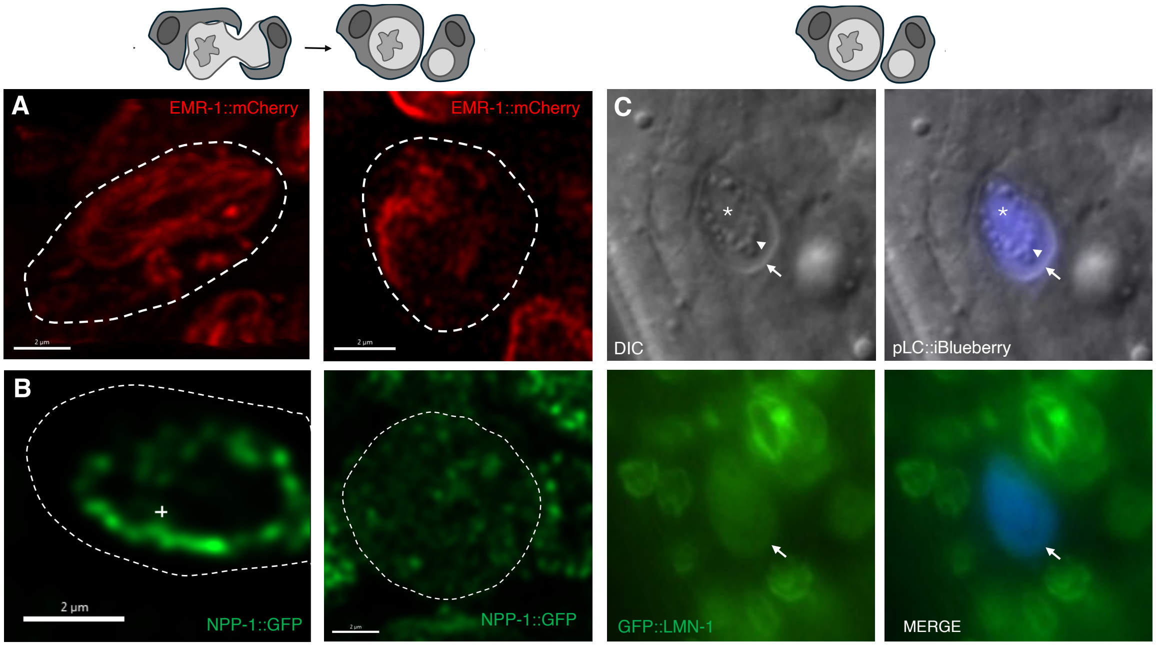


**Supplemental Figure 1. Nuclear envelope protein dynamics during cell splitting.**
(A) Confocal imaging of EMR-1::mCherry during competitive phagocytosis (left) and split (right) stages. Scale bar, 2µm.
(B) Confocal imaging of NPP-1::GFP during competitive phagocytosis (left) and split stages (right). Scale bar, 2µm. + indicates nucleoporin signal within the nuclear envelope membrane.
(C) Split linker cells exhibit perinuclear blebbing (arrowhead in DIC) and diffuse lamina (arrow). Asterisk marks the nucleus.

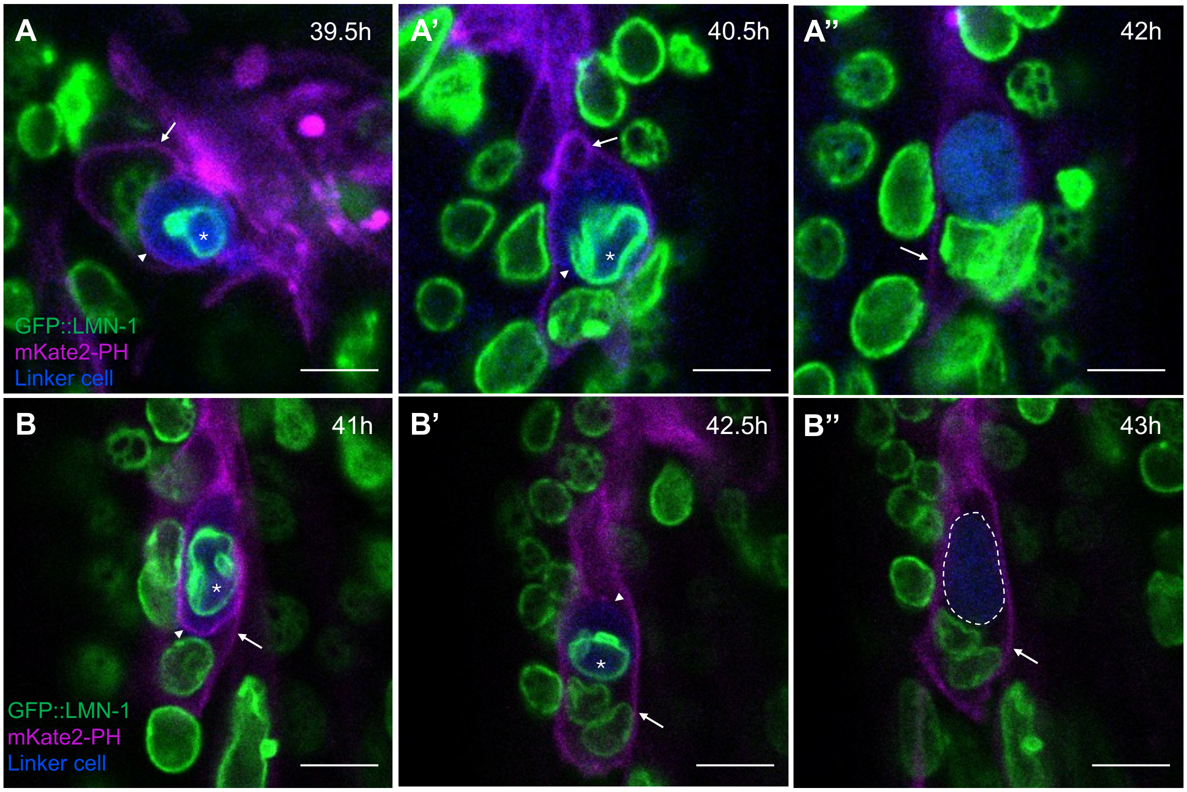


**Supplemental Fig 2. Lamina disassembly occurs after phagosome maturation.**

(A-B") Confocal live-imaging of a linker cell labeled with *mig-24*p::iBlueberry, lamin endogenously tagged with GFP, and mKate2-PH labeling unsealed phagosome membranes and plasma membrane of the engulfing cells. Asterisk, intact lamina of the linker cell nucleus. Arrowhead, unsealed phagosome membrane. Arrow, engulfing cell membrane. (A-A'') – animal 1, scale bar, 4μm; (B-B")'– animal 2, scale bar, 5μm

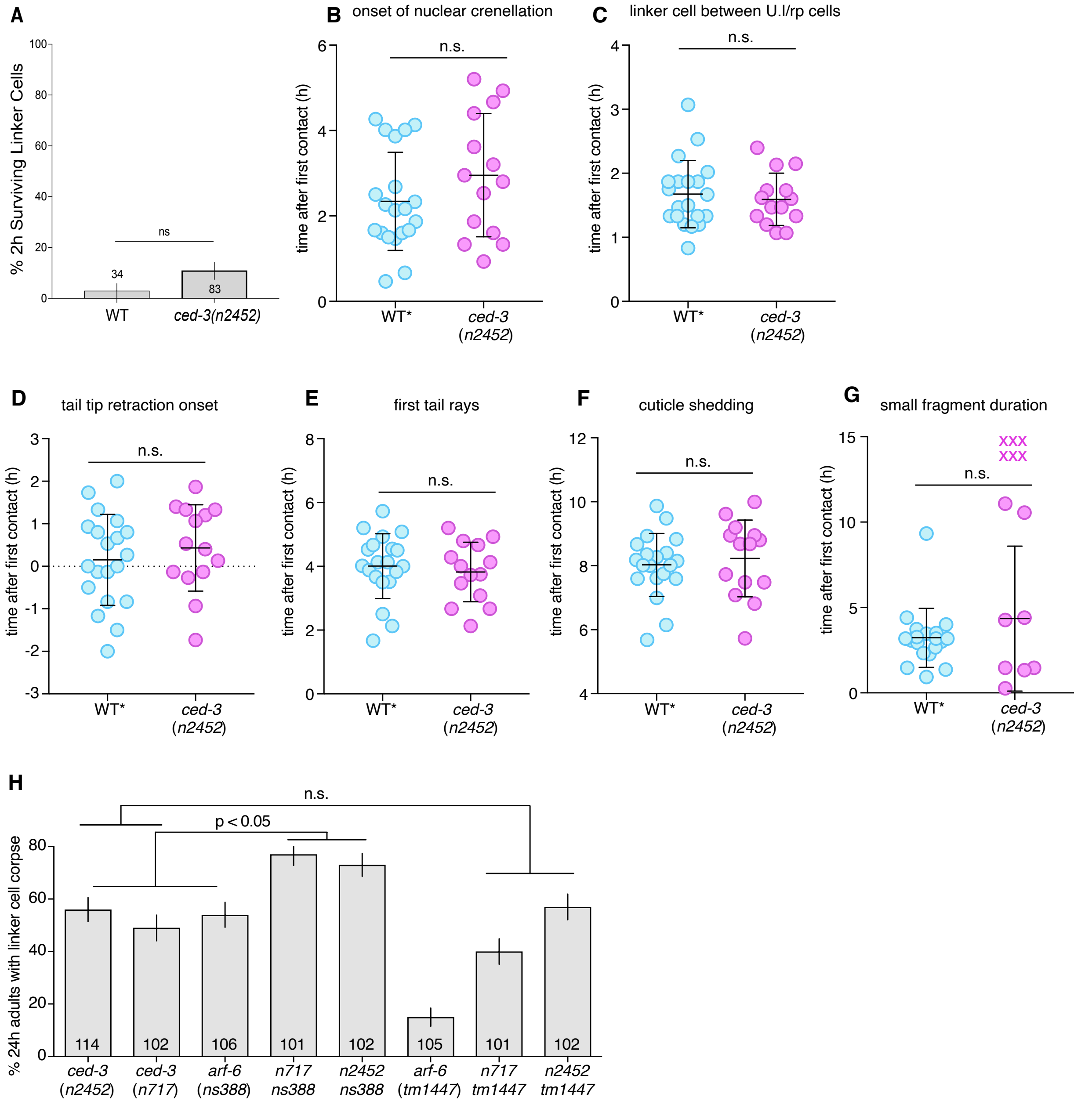


**Supplemental Figure 3. *ced-3* and *arf-6* genetically interact.**
(A) ced-3(n2452) 2h adults have normal linker cell death initiation similar to wild-type. The number of animals scored inside bars. Error bars, standard error of sample proportion, Fisher's exact test. ns is p>0.05.
(B-G) Developmental milestones in *ced-3(n2452)* animals. Wild-type data taken from [16]. n.s., not significant. Student's T test.
(H) Linker cell degradation in indicated genotypes. Strains contain *lag-2p::GFP* linker cell reporter and *him-5(e1490)*. Number of animals scored inside bars. Error bars, standard error of the proportion. Fisher's exact test.

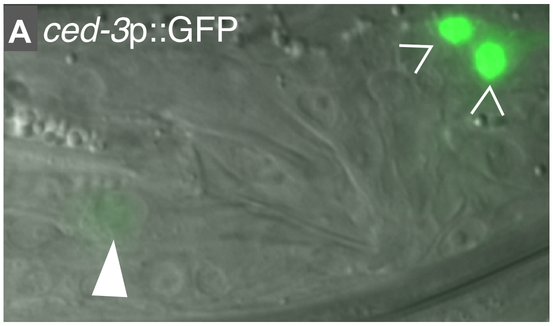


**Supplemental Figure 4. *ced-3* expression is lower in the linker cell than in apoptotic cells.**
(A) Transgene *ced-3*p::GFP is expressed in the dying linker cell (triangle) and in the cells dying by apoptosis (arrowheads).

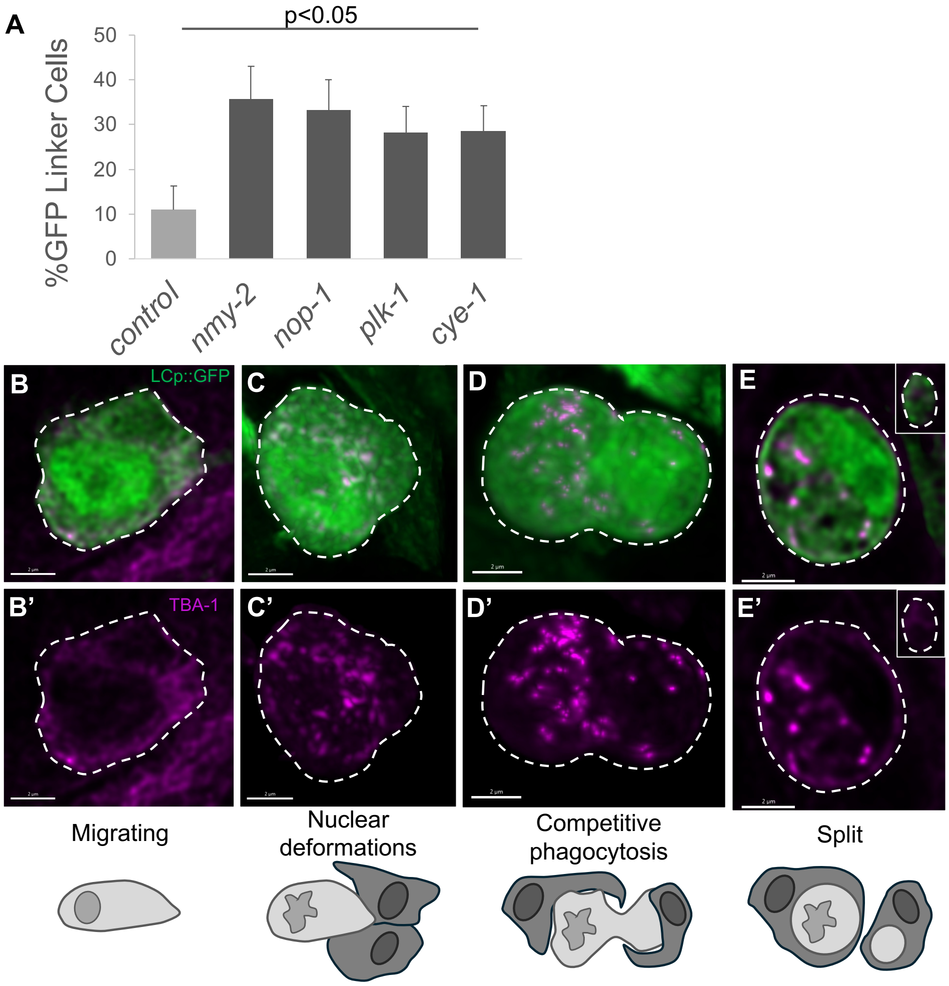


**Supplemental Figure 5. Cell cycle genes in LCD.**
(A) Linker cell-specific RNAi against cytokinesis (*nmy-2*, *nop-1*) and cell cycle (*plk-1*, *cye-1*) genes. Animals were scored for linker cell persistence at 8h post-molt. Error bars, standard error of the proportion. Fisher's exact test.
(B-E') Confocal imaging of Tubulin/TBA-1 (TBA-1::mCherry) in the linker cells marked by *lag-2p*::GFP (LCp) during migration (before death onset), and during cell death progression at the stages of nuclear deformations, competitive phagocytosis and split. Scale bar, 2μm.

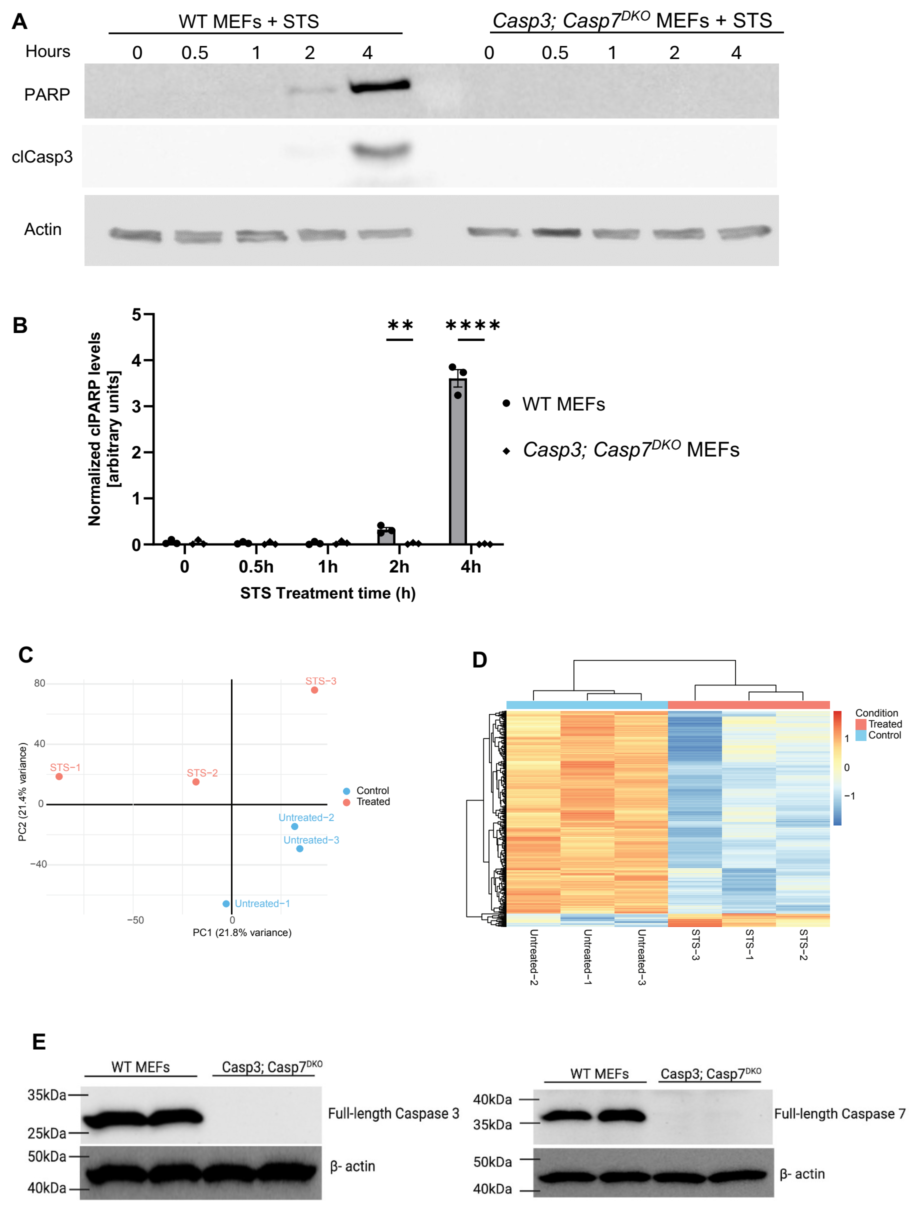


**Supplemental Figure 6. Additional MEFs data and RNA-seq quality control.**
(A) SDS-PAGE and Western Blot analysis of wild-type and *Casp3; Casp7*$^{DKO}$ MEFs treated with 1.5 μM STS for 0.5, 1, 2, and 4 hours and probed with antibodies against cleaved PARP and cleaved Caspase3. Actin is a loading control.
(B) Densitometry analysis of cleaved PARP levels. Three biologically independent Western blot experiments.
(C) Principal component analysis of DMSO-treated and STS-treated wild-type MEFs.
(D) Unsupervised hierarchical clustering of DMSO-treated (blue, control) and STS-treated (red, treated) wild-type MEFs.
(E) Western blot analysis of *Casp3; Casp7*$^{DKO}$ MEFs to validate the genotype. Blots were probed with antibodies against Caspase 3 and Caspase 7. Beta-actin is a loading control.
